# Spatially Patterned Bi-electrode Epiretinal Stimulation for Axon Avoidance at Cellular Resolution

**DOI:** 10.1101/2021.08.17.454395

**Authors:** Ramandeep S. Vilkhu, Sasidhar S. Madugula, Lauren E. Grosberg, Alex R. Gogliettino, Pawel Hottowy, Wladyslaw Dabrowski, Alexander Sher, Alan M. Litke, Subhasish Mitra, E.J. Chichilnisky

## Abstract

**Objective:** Epiretinal prostheses are designed to restore vision to people blinded by photoreceptor degenerative diseases by stimulating surviving retinal ganglion cells (RGCs), which carry visual signals to the brain. However, inadvertent stimulation of RGCs at their axons can result in non-focal visual percepts, limiting the quality of artificial vision. Theoretical work has suggested that axon activation can be avoided with current stimulation designed to minimize the second spatial derivative of the induced extracellular voltage along the axon. However, this approach has not been verified experimentally at the resolution of single cells. *Approach*. In this work, a custom multi-electrode array (512 electrodes, 10 μm diameter, 60 μm pitch) was used to stimulate and record RGCs in macaque retina *ex vivo* at single-cell, single-spike resolution. RGC activation thresholds resulting from bi-electrode stimulation, which consisted of bipolar currents simultaneously delivered through two electrodes straddling an axon, were compared to activation thresholds from traditional single-electrode stimulation.

**Results:** On average, across three retinal preparations, the bi-electrode stimulation strategy reduced somatic activation thresholds (∼21%) while increasing axonal activation thresholds (∼14%), thus favoring selective somatic activation. Furthermore, individual examples revealed rescued selective activation of somas that was not possible with any individual electrode. *Significance*. This work suggests that a bi-electrode epiretinal stimulation strategy can reduce inadvertent axonal activation at cellular resolution, for high-fidelity artificial vision.

**Novelty & Significance:** The effectiveness of bi-electrode stimulation for enhancing the electrical activation of retinal neurons was tested using high-density multi-electrode recording and stimulation in isolated macaque retina. The results suggest that spatially patterned bi-electrode stimuli reduce unwanted axon activation and thus improve the selectivity of stimulation at cellular resolution. Similar patterns could be implemented in a future high-resolution prosthesis to permit a more faithful replication of normal retinal activity, at the resolution of single-cells and single-spikes, for the treatment of incurable blindness.

## Introduction

Epiretinal prostheses are designed to restore visual function in people blinded by photoreceptor degenerative diseases such as age-related macular degeneration or retinitis pigmentosa [1,2]. These devices are implanted on the retinal ganglion cell (RGC) side of the retina, and electrically activate RGCs that have survived the degeneration process, causing them to transmit artificial visual signals to the brain. Although devices implanted in patients have demonstrated some rudimentary vision restoration, the reported visual acuity and clinical value provided by existing devices is low [3–7].

One major problem limiting the visual acuity from existing epiretinal devices is the inadvertent activation of RGC axons [8,9]. In particular, activation of axons that originate from RGCs with visual receptive fields spatially distant from the location of electrical stimulation causes the perception of non-focal elongated phosphenes, limiting the quality of artificial vision [10].

In principle, selective activation of RGC somas while avoiding axons is possible with the use of effective temporal and spatial stimulation strategies, and several approaches have been proposed. Temporal stimulation strategies, which involve modulations of the stimulus waveform leading to maximally differentiable axon and somatic activations, include low frequency stimuli [11], cathodal monophasic pulsed stimuli [12], and charge-balanced long-duration bi-phasic pulsed stimuli [8]. Alternatively, spatial stimulation strategies leverage spatial biophysical activation characteristics of RGC axons to minimize axonal activation [13,14]. Examples of spatial strategies include rectangular electrodes positioned to straddle an axon [15,16] and patterned multi-electrode stimulation on a dense multi-electrode array [17].

While such approaches improve selective activation of RGC somas over axons to a degree, they fail to activate RGCs at the natural spatiotemporal resolution of the retina’s neural code – single cells and single spikes. Specifically, a challenge for high-resolution epiretinal stimulation is the fact that different types of RGCs, each encoding unique aspects of the visual scene, are intermixed on the surface of the retina. Each RGC type signals visual information with precisely timed spikes that are highly stereotyped in all cells of each type. Hence, focal stimulation at the resolution of single cells and single spikes is likely required to reproduce high-fidelity vision. The temporal stimulation strategies discussed above depend on stimulus waveform modulations leading to network-mediated RGC activation, as opposed to direct RGC activation, and thus fail to match the temporal resolution of direct RGC activation (but see [18]). In contrast, the spatial stimulation strategies result in direct RGC activation, but the studies were either based in simulation or used experimental setups that failed to provide cellular and cell type resolution. Hence, it remains unclear whether such strategies can match the spatial resolution of the retina for use in a high-resolution epiretinal device.

Here, we test whether a biophysically-inspired spatial bi-electrode strategy can improve selectivity for stimulating RGC somas over axons at cellular resolution. Using large-scale, high-density recording and stimulation in isolated macaque retina [19–22], we evaluate how effectively a given target RGC soma can be activated without activating a passing non-target axon, with both bi-electrode and single-electrode stimulation. In many cases across three retinal preparations, the bi-electrode strategy enhanced the selective activation of target RGC somas over non-target passing RGC axons. Furthermore, individual examples reveal rescued selective activation of somas that was not possible with any individual electrode, confirming the potential value of the approach for artificial vision.

## Methods

### Experimental setup

A custom 512-electrode system [19,22,23] was used to stimulate and record from RGCs in three isolated rhesus macaque monkey (*Macaca mulatta*) retinas. Retinas were obtained from terminally anesthetized animals euthanized during the course of research performed by other laboratories. All procedures were performed in accordance with institutional and national guidelines and regulations. Briefly, eyes were hemisected in room light following enucleation. Next, the vitreous was removed and the posterior portion of the eye containing the retina was kept in darkness in warm, oxygenated, bicarbonate buffered Ames’ solution (Sigma). Patches of retina ∼3 mm on a side were isolated under infrared light, placed RGC side down on the multielectrode array, and superfused with Ames solution at 35°C. Electrodes were 8-15 μm in diameter and arranged in a 16 × 32 isosceles triangular lattice with 60 μm spacing between adjacent electrodes [24]. Within an experiment, platinization of the electrodes produced relatively uniform noise (and thus effective electrode sizes), e.g. ±6% (standard deviation) across electrodes. A platinum wire encircling the recording chamber (∼1cm diameter) served as the distant return electrode. Voltage recordings were band-pass filtered between 43 and 5,000 Hz and sampled at 20 kHz. Spikes from individual RGCs in the voltage recordings were identified and sorted using standard spike sorting techniques [24].

### Visual stimulation and cell type classification

To identify the type of each cell recorded, the retina was visually stimulated by a dynamic white noise stimulus, and the spike-triggered average (STA) stimulus was computed for each RGC, as previously described [25,26]. The STA summarizes the spatial, temporal, and chromatic structure of light response. In particular, clustering on the spatial (receptive field size) and temporal (time course) components of the STA was performed to identify distinct cell types, as previously described [27]. Analysis focused on ON and OFF parasol RGCs due to the high SNR of their recorded spikes, which was useful for reliable spike sorting in the presence of electrical artifacts (see below).

### Electrical image

The electrical image (EI) represents the average spatiotemporal pattern of voltage deflections produced on each electrode of the array during a spike from a given cell [24]. Electrical images were calculated from data recorded during visual stimulation and served as spatiotemporal templates for the spike waveforms of the cells to be detected during electrical stimulation.

The spatial positions of relevant cell compartments (axon and soma) of a recorded cell relative to the electrode array were estimated using the electrical image. For each recorded cell, the shape of the spike waveform from the EI on an electrode was used for compartment identification: a triphasic waveform was taken to indicate recording from the cell axon whereas a biphasic waveform was taken to indicate recording from the cell soma [24]. Then, the temporal component of the EI was collapsed by computing the maximum negative amplitude over all electrodes. Using this purely spatial EI representation, a two-dimensional Gaussian was fitted over all electrodes recording somatic signals to estimate the geometric area on the array electrically coupled to the soma, and a quadratic polynomial was fitted to all electrodes recording axonal signals to estimate the axon trajectory (Fig. 1A) [28].

**Figure 1.**
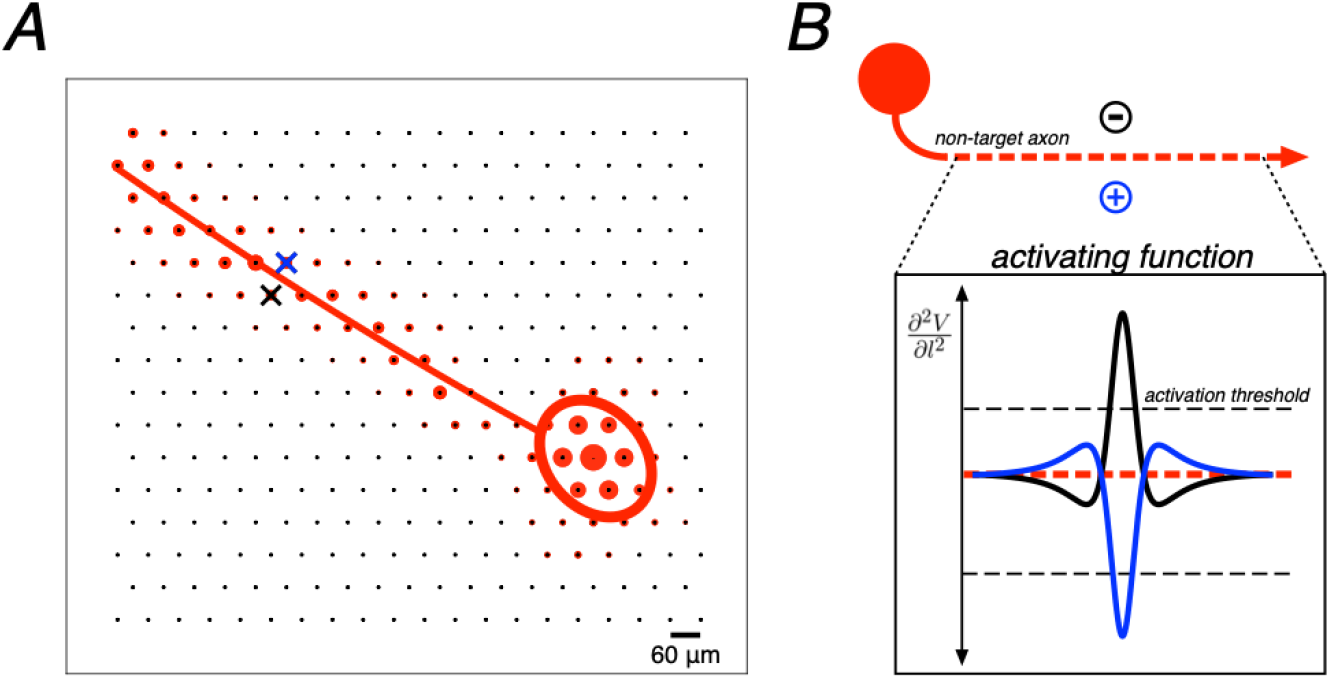
Example electrical image decomposition into axonal and somatic compartments along with biophysical theory. A. Spatial component of the electrical image for a single cell plotted on the electrode array along with polynomial axon trajectory fit and Gaussian somatic compartment fit. Crosses represent an example bi-electrode stimulation pattern, straddling the axon fit (blue cross, positive polarity electrode; black cross, negative polarity electrode). B. Example of an ideal bi-electrode stimulus: a pair of electrodes perfectly straddling an axon providing equal and opposite currents. The resulting activating function (the second spatial derivative of voltage along the length of the axon) from each active electrode is shown. Linear summation of electric fields implies that these activating functions would cancel perfectly in this idealized case.

### Electrical stimulation

Electrical stimulation was provided through one or more electrodes while recording RGC activity from all electrodes. Two types of stimulation patterns were tested: single-electrode stimulation and bi-electrode stimulation. The single-electrode stimulation consisted of a charge-balanced, triphasic pulse passed through one electrode. The triphasic pulse was composed of anodal/cathodal/anodal phases with relative current amplitudes 2:-3:1 and phase duration of 50 μs per phase (150 μs total). These parameters were chosen to minimize the electrical artifact [19,20,22]. Single-electrode stimulation was delivered in 25 repeated trials at 40 current amplitudes (10% increments) between 0.1 and 3.7 μA. The bi-electrode stimulation consisted of the same single-electrode current pattern passed through one electrode, but with equal and opposite polarity current simultaneously passed through an adjacent electrode. The stimulating electrodes were chosen based on highest SNR of recorded spikes and the electrodes were spatially patterned such that the pair of simultaneously active electrodes straddled a passing axon, as estimated by the fitted axon trajectory (see above), to minimize the activating function along the length of the axon [13] (Fig. 1B). Bi-electrode stimulation was supplied for multiple trials at varying current amplitudes (10% increments) between 0.2 and 3.8 μA (retina 1; 40 repetitions), 0.2 and 4.9 μA (retina 2; 20 repetitions), or 0.2 and 4.3 μA (retina 3; 25 repetitions). The ordering of the stimulation patterns was chosen pseudo-randomly, restricted so that each successive stimulating electrode or pair of electrodes was far from the previous and subsequent stimulating electrodes. This was done to avoid stimulating the same cell(s) in rapid succession.

### Responses to electrical stimulation

The spikes recorded during electrical stimulation were analyzed using a custom supervised template matching method [20]. First, the electrical image of each cell was calculated from visual stimulation data, and served as a template for the spike waveform of each cell to be detected during electrical stimulation. In the electrical stimulation data, an automated algorithm separated spikes from the electrical artifact by grouping traces according to the artifact waveform estimate and the spike waveform of each cell analyzed. The resulting electrically elicited spike waveforms were visually inspected for sorting errors and manually corrected as needed.

For each analyzed cell, spike probabilities were calculated across 20-40 trials at each stimulus current amplitude, and cell spiking probability was modeled as a function of current amplitude by a sigmoidal relationship (Equation 1).

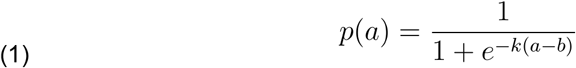

where a is the current amplitude, p(a) is the spike probability of a given cell, and k and b are free parameters (representing sigmoidal slope and threshold, respectively). Fitted sigmoidal curves were used to compute the *activation threshold*, defined as the current amplitude producing 50 percent spiking probability.

### Statistical analysis of threshold and selectivity changes

The relationship between activation thresholds obtained from bi-electrode and single electrode stimulation was analyzed using a resampling approach. To determine the estimated variation in the slopes of lines fitted in Fig. 2 and Fig. 3, data were resampled with replacement, producing simulated data sets with the same number of points as the real data set, drawn from the distribution given by the measured data. This resampling was repeated 1000 times and, for each iteration of resampling, the slope of the least squares linear fit to the data was computed. Values in the text correspond to the slope values obtained from the measured data along with 90% confidence intervals based on the resampled data.

**Figure 2.**
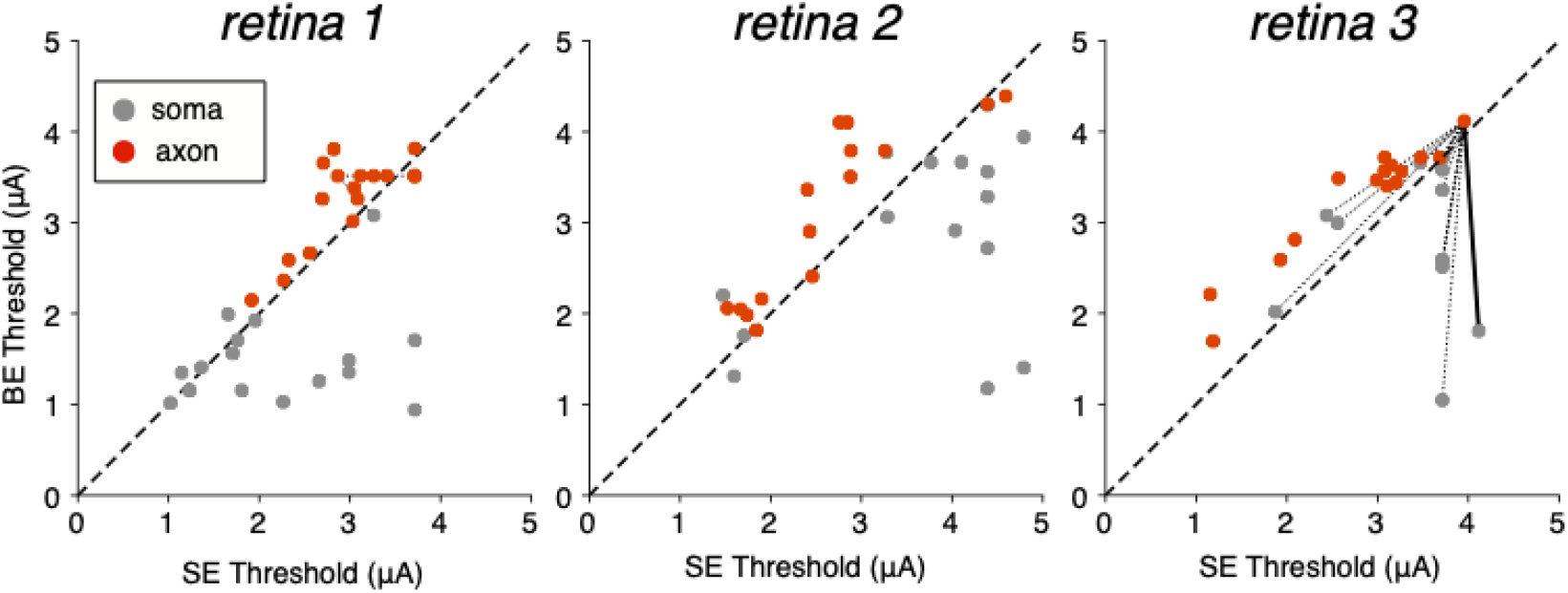
Axonal and somatic activation thresholds from bi-electrode and single-electrode stimulation. Scatter plots for three preparations show activation threshold of analyzed cells to bi-electrode (BE) and single-electrode (SE) stimulation. Dashed diagonals denote equality. An example of the “surrogate” analysis method (see Results) is illustrated in retina 3: dashed lines show all possible pairings between a non-target axon and all somas; solid line denotes a soma-axon pair targeted by the same set of electrodes. Data are shown from 40 distinct cells and 192 distinct cell-electrode pairs across the three retinal preparations.

**Figure 3.**
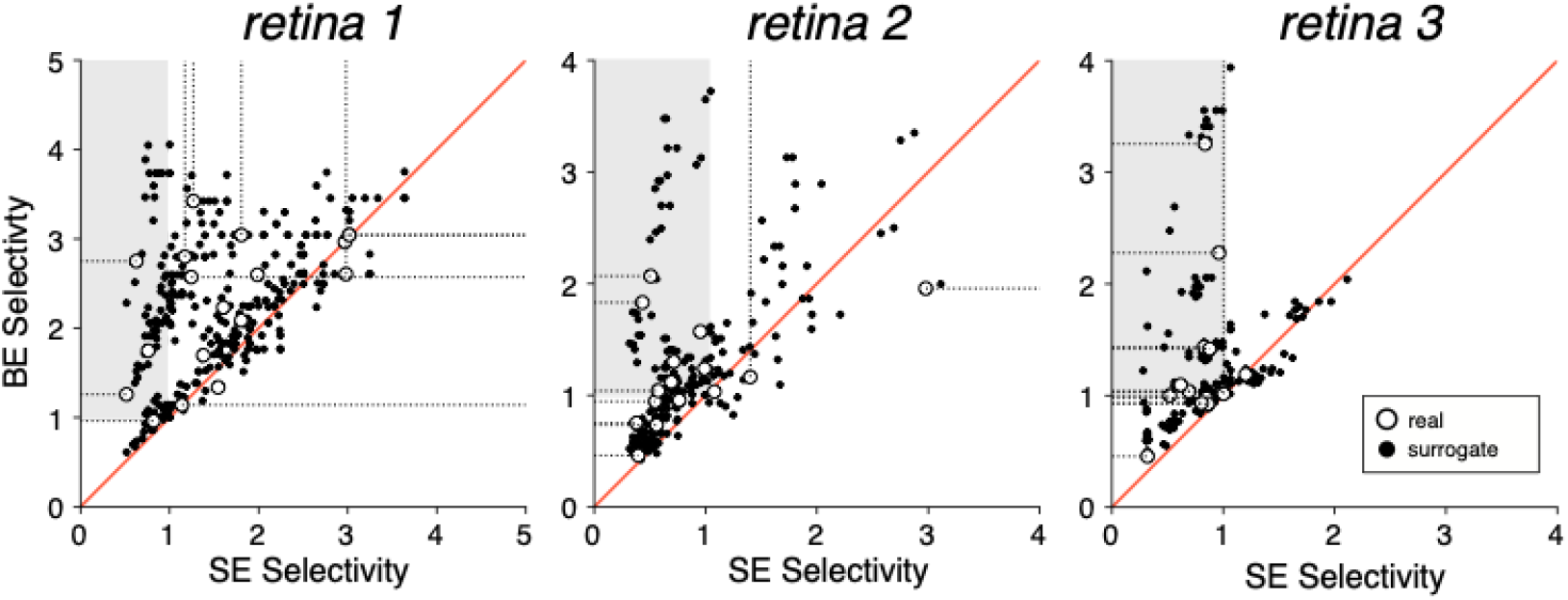
Impact of bi-electrode stimulation on selectivity. Scatter plots with each point representing selectivity calculated for a target soma (gray points, Fig. 2) and non-target axon (red points, Fig. 2) pair with bi-electrode (BE) and single-electrode (SE) stimulation in three retinas. Red diagonals denote equality. Shaded region represents regions of rescued selectivity (Fig. 4). Open symbols with dotted tails represent scenarios in which the target soma or non-target axon did not spike at the maximum supplied current amplitude. In these cases selectivity was calculated with the threshold set to the largest current amplitude tested, and the tail shows the possible range of true selectivity for larger thresholds.

## Results

Electrical recording and stimulation were performed on isolated macaque retinas using a high-density 512-electrode array (see Methods). Direct RGC activation caused by current pulses passed through one or two electrodes simultaneously was investigated based on biophysically derived insight into the mechanism of axonal activation [14,29]. To determine the effectiveness of bi-electrode stimulation to selectively activate cells, stimulation of a target soma and a non-target axon with a single electrode on one side of the axon was compared to bi-electrode stimulation with an equal and opposite current pulse simultaneously through a second electrode on the opposite side of the axon.

After delivering 20-40 current pulses (triphasic, charge-balanced, 50 μs per phase, 150 μs total duration) at 30-40 current levels, a sigmoidal relationship was used to characterize the probability of RGC activation as a function of injected current. The current amplitude level corresponding to RGC activation on 50% of stimulation trials was estimated and defined as the *activation threshold* (see Methods). Stimulation was performed near the soma of a target RGC and near the axon of a non-target RGC using information from the recorded electrical images (EIs) of the cells. The EI represents the average spatiotemporal pattern of voltage deflections produced on each electrode of the array during a spike from a given cell (see Methods). Somatic or axonal stimulation was achieved by using the EIs to determine electrodes that recorded unambiguous somatic (biphasic) or axonal (triphasic) spikes from the target RGC soma or the non-target RGC axon, respectively [24]. To determine the position and orientation of the axon with respect to the stimulating electrodes, a quadratic spline was fitted to interpolate the trajectory of the axonal compartment of the electrical image [28].

### Effect of bi-electrode strategy on somatic and axonal RGC activation thresholds

Comparison of thresholds obtained with single-electrode and bi-electrode stimulation revealed two trends. First, as predicted, bi-electrode stimulation systematically increased axonal thresholds (Fig. 2). Second, on average, bi-electrode stimulation decreased somatic thresholds, however, there was noticeable variability in the observed threshold changes (see Discussion) (Fig. 2). To quantify the relationship between bi-electrode and single-electrode stimulation of axons and somas across retinas, the slopes of the relation between bi-electrode and single-electrode activation thresholds (Fig. 2) were computed, along with 90% confidence intervals on the slope obtained by resampling (see Methods; Table 1). For cells activated at somatic compartments (grey points, Fig. 2), on average, bi-electrode stimulation resulted in lower thresholds than single-electrode stimulation (slopes below 1) in all three retinas. In contrast, for cells activated at axonal compartments (red points, Fig. 2), bi-electrode stimulation produced higher thresholds compared to single-electrode stimulation in all three retinas. These overall trends suggest that bi-electrode stimulation could be used to selectively activate somas over axons.

**Table 1.**
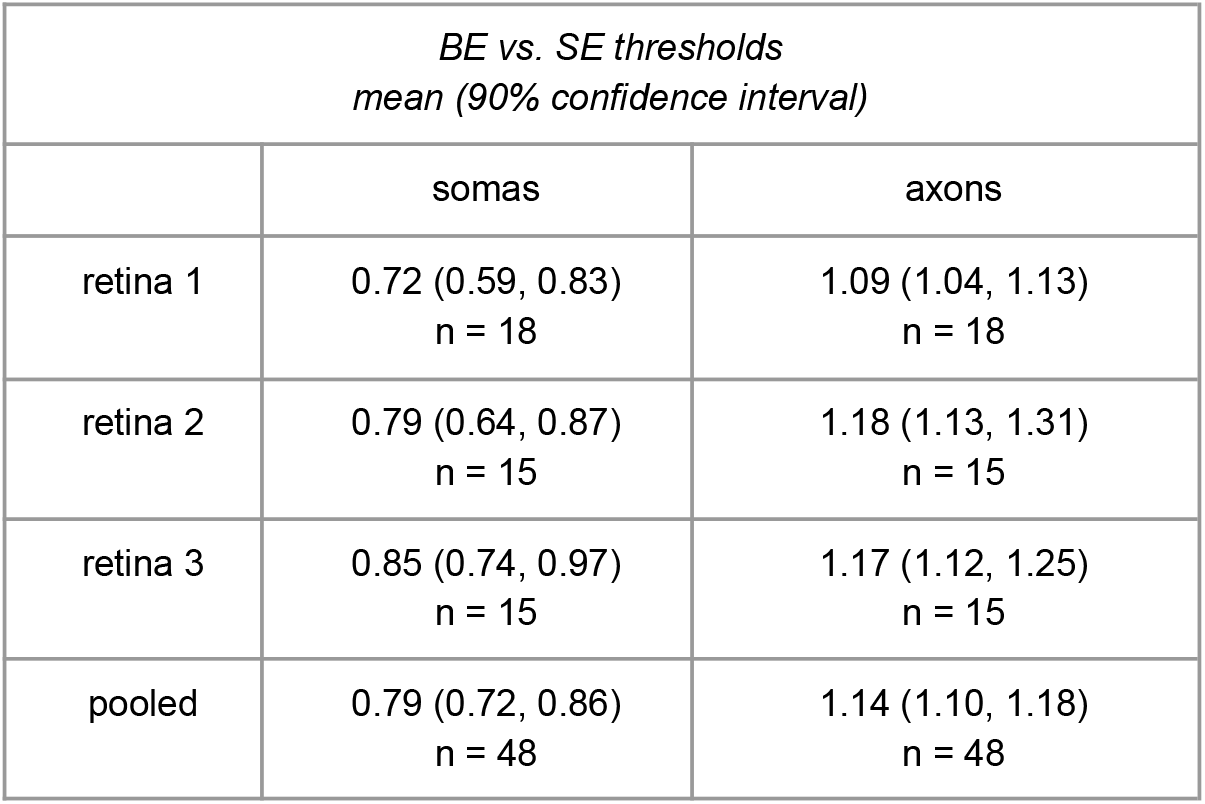
Change in axon and soma activation thresholds with bi-electrode vs single-electrode stimulation. Columns of the table represent RGCs activated at somatic compartments versus RGCs activated at axonal compartments (somas and axons). Each entry indicates the fitted slope of the data in Fig. 2, with 90% confidence intervals on the slope obtained by resampling with replacement 1000 times (see Methods). The last row shows the aggregate statistics from all cells across all three retinas.

### Bi-electrode strategy enhances selective activation of target soma over non-target axon

It is unclear from the thresholds alone how useful the strategy will be in real scenarios of a single axon passing by a target soma. To test the usefulness of the approach, specific instances of selective activation of target somas over nearby non-target axons were examined. The target soma and non-target axon were stimulated by the same electrode or pair of electrodes using single-electrode or bi-electrode stimulation, respectively. *Selectivity*, defined as the ratio of non-target axon activation threshold to target soma activation threshold, was computed for both bi-electrode and single-electrode stimulation. The selectivity for activating a target soma over a non-target axon was systematically higher for bi-electrode stimulation versus single-electrode stimulation in all three retinas (Fig. 3, open symbols).

In the above analysis, the number of soma-axon pairs stimulated by the same pair of electrodes was limited by the difficulty of reliably spike sorting the cell response to electrical stimulation. Therefore, a surrogate analysis was also performed, comparing the activation thresholds of all pairs of somas and axons in the same retina (Fig. 2, surrogate pairing visual). In this analysis, the target soma and non-target axon were in fact stimulated by different electrodes on the array, but the increased number of analyzable pairs served to increase the statistical power of the single-electrode vs. bi-electrode comparison. Selectivity was systematically higher with bi-electrode stimulation (Fig. 3, closed symbols). Slopes fitted to these data, along with 90% confidence intervals on the slope obtained by resampling (see Methods), confirmed the trend in all three retinas and in the data pooled across retinas (Table 2). These results were consistent with results from true soma-axon pairings (above), demonstrating the robustness of the trends to potential sources of variability across electrodes and cells.

**Table 2.** Change in selectivity with bi-electrode vs single-electrode stimulation. Each entry indicates the fitted slope of the bi-electrode versus single electrode selectivity data from Fig. 3, with 90% confidence intervals on the slope obtained by resampling from the data with replacement 1000 times (see Methods). The last row indicates the aggregate results from all cells across all three retinas.

| *Selectivity measured by thresholds*<br>*mean (90% confidence interval)* |  |  |
| --- | --- | --- |
|  | bi-electrode vs single electrode<br>(surrogate pairings) | bi-electrode vs single electrode<br>(real pairings) |
| retina 1 | 1.44 (1.37, 1.49) | 1.39 |
| retina 2 | 1.55 (1.44, 1.64) | 1.40 |
| retina 3 | 1.48 (1.40, 1.57) | 1.55 |
| pooled | 1.47 (1.42, 1.51) | 1.44 |

### Rescued selectivity

In some cases, bi-electrode stimulation made it possible to selectively activate a soma that could not be selectively activated with any nearby individual electrode (three examples shown in Fig. 4). Specifically, in many soma-axon pairs recorded on the same electrodes (n = 29), it was not possible to selectively activate the soma without unwanted axonal activation of a distant cell using the electrodes near the cell (SE selectivity < 1; Fig. 3). However, in a majority of cases (18/29), bi-electrode stimulation resulted in selective activation of the soma without activation of the axon, rescuing selectivity (BE selectivity > 1; Fig. 3 shaded region).

**Figure 4.**
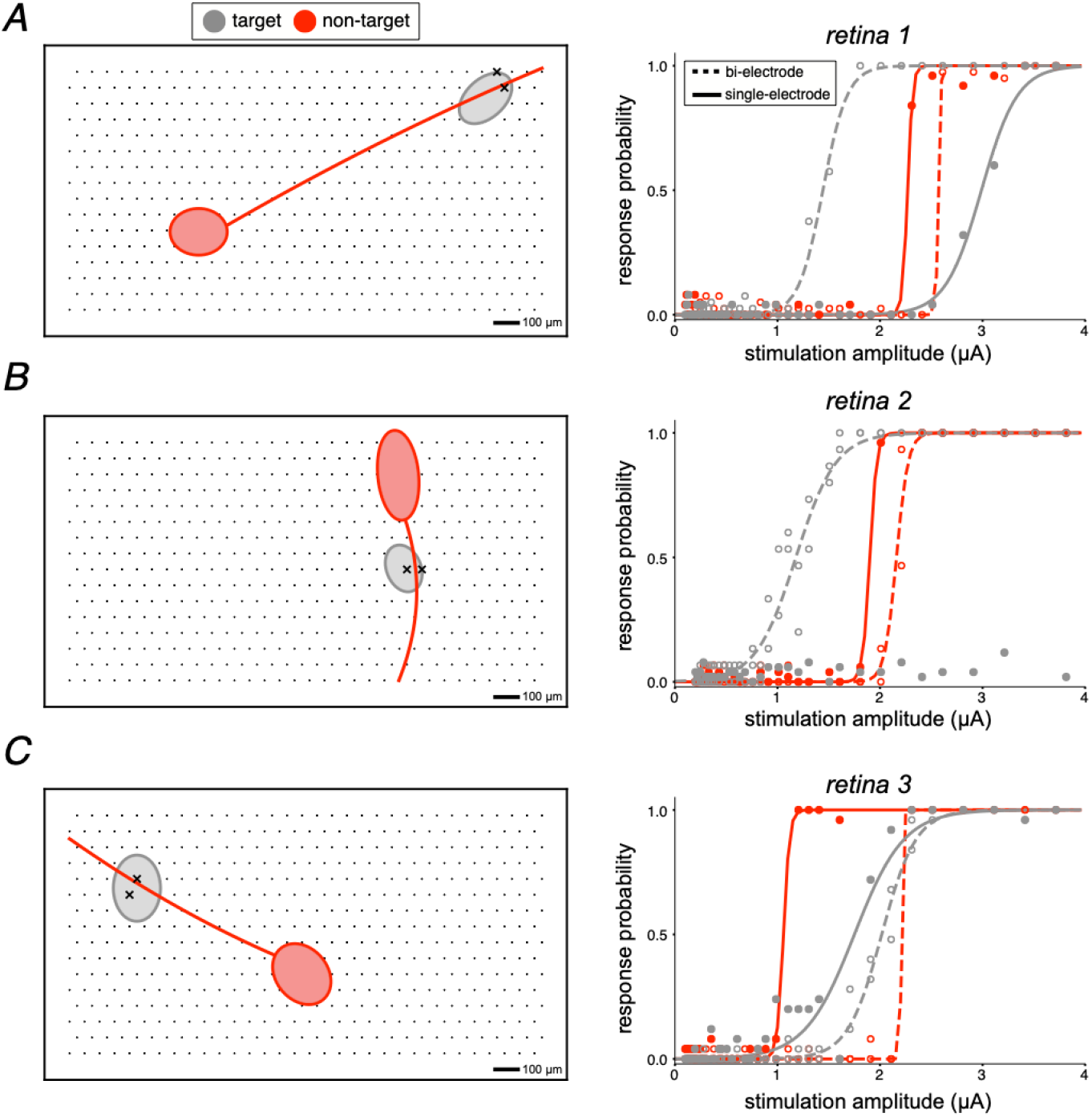
Electrical images and activation curves for target somas and non-target axons exhibiting rescued selectivity. Each panel on the left shows a simplified electrical image of the axonal and somatic compartments of a recorded cell over a region of the electrode array. Each panel on the right shows the observed activation curves for the target and non-target cell, using bi-electrode and single electrode stimulation. In all cases, bi-electrode stimulation resulted in selective activation of the target soma, at a lower current level than the threshold for the non-target axon, which was not possible with single electrode stimulation. Soma-axon pairs A, B, and C were from retina 1, 2, and 3, respectively.

## Discussion

The results indicate that bi-electrode stimulation can enhance the selectivity of stimulating RGC somas over axons in the primate retina (Fig. 3). Furthermore, bi-electrode selectivity improvements were demonstrated by individual scenarios in which a specific soma could be effectively activated without activation of a specific nearby axon, in a way that was not possible with the individual electrodes of the array (Fig. 4). This is significant because activation of a distant cell, encoding a distant region of the visual field, at its axon can cause unwanted visual information to be transmitted to the brain [10]. These findings are consistent with previous simulation and experimental work [15–17]. The primary novel aspect of the present study is that RGC activation was probed at the natural spatiotemporal resolution of the retinal code -- single cells and single spikes -- which may be important for high-fidelity artificial vision with future clinical implants.

While no previous work has explored axon avoidance at cellular resolution using large-scale recordings, several other strategies have been proposed and tested. These can be broadly divided into spatial and temporal approaches.

One spatial approach used stimulation with long rectangular electrodes positioned along an axon. This strategy was found to minimize axon activation in simulation and experiments [15,16]. Such spatial strategies leverage biophysical activation characteristics of axons [14,29], and the best-case experimental results show a 34% difference between distant cell and local cell activation thresholds, presumably reflecting axon avoidance. However, proper orientation of the rectangular electrodes present a challenge during surgical implantation. Moreover, the geometry of these electrodes limits the ability to reliably target a single cell and the study found that the spatial resolution of stimulation was not improved by simply reducing the size of the electrodes (see [30]). Another study simulated axon avoidance using dense multi-electrode arrays and simultaneous stimulation at multiple smaller electrodes, mimicking stimulation with long electrodes [17]. However, the study was limited to computational simulations and the approach was not tested experimentally.

Temporal modulations of the current stimulus have also been found to effectively reduce unwanted axonal activation. These strategies predominantly use long-duration current pulses (>25 ms) [8] or low frequency (<25 Hz) sinusoidal stimulation [11,31], and can produce ∼3x selectivity improvements. However, long pulse durations also tend to activate RGCs indirectly through the retinal network, typically producing multiple spikes with longer and more variable latencies, thereby making it difficult to match the natural temporal resolution of the retina. A different approach [18] relied on short-duration pulsed stimuli (<0.10 ms), which produce direct RGC activation, and demonstrated ∼3x selectivity improvement for activating somas over axons. Another study [32] corroborated the use of short-duration pulses (<0.12 ms), and also explored the design of asymmetric anodic-first pulses to further enhance selective activation of somas. While these studies showed significant selectivity improvements, larger-scale studies performed with many cells and retinas have demonstrated that short-duration stimulation leads to significant inadvertent axon activation [22,33,34]. Therefore, it is unclear whether such temporal modulation strategies alone will improve the performance of clinical devices.

The proposed bi-electrode strategy can be viewed as a hybrid of previous spatial and temporal approaches. The spatial arrangement of the stimulating and return electrode straddling the axon exploited biophysical properties of axon activation, and the short stimulation pulses produced direct activation, resulting in a 21% reduction in somatic activation thresholds and 14% increase in axonal activation thresholds -- a 1.44x selectivity improvement. Furthermore, the performance of this bi-electrode strategy was measured *ex vivo* with hardware and conditions intended to mimic the function of a future clinical device.

Although axonal thresholds increased with bi-electrode stimulation, somatic thresholds sometimes increased and sometimes decreased (Fig 2). A possible explanation is that the secondary electrode deflects current flow in a direction either towards or away from excitable cell structures (e.g., the axon initial segment) [20,35–38] depending on the geometry of the electrodes relative to the cell. A related possibility is that current is deflected toward or away from multiple activation sites, non-linearly modifying the probability of activation [39]. While these hypotheses deserve further examination, the overall trend was that bi-electrode stimulation enhanced selectivity significantly in spite of somatic threshold variability.

The cell types examined in this study were ON and OFF parasol cells, two numerically dominant cell types in primates (∼16% of the RGC population [40], in the peripheral retina. A major reason for this focus is that the large spikes produced by these cells were comparatively easy to identify in the presence of the electrical stimulation artifact. However, other cell types (e.g., midget cells: comprising ∼50% of the RGC population) must be investigated in the future for a more comprehensive study of the efficacy of bi-electrode stimulation. Furthermore, future devices target implantation in more central regions of the retina (e.g., the raphe) to enable the highest resolution artificial vision, and the bi-electrode strategy has yet to be investigated in these regions. Additionally, the analysis focused on the relative threshold changes of a single target cell and single non-target cell, ignoring other cells near the stimulation site(s). In reality, threshold changes in all nearby cells must be accounted for to comprehensively determine the selectivity improvement from bi-electrode stimulation. However, it was difficult to investigate threshold changes for many non-target cells because the process of manually sorting spikes after electrical stimulation is time-consuming [20,22]. One potential solution would be the development of algorithms to automate spike sorting in the presence of electrical artifacts, particularly for multi-electrode stimulation, an effort that is underway [41].

The biophysical motivation of the bi-electrode strategy is that it decreases the magnitude of the induced electric field along the longitudinal direction of an axon, resulting in decreased axon activation. However, the electric field near the stimulating electrodes will vary with different stimulation hardware geometries. The present work tested this strategy on a 60 μm pitch electrode array with ∼10 μm diameter electrodes. Variations in electrode size, pitch, and shape would likely result in variations of the induced electric field along the axon, and thus variation in activation thresholds. Although exploration of novel electrodes could further improve axon suppression through tailored electric field shaping, such changes could impact the ability to focally stimulate cells. For example, long rectangular electrodes have been shown to shape electric fields in a manner that minimizes axon activation (see above; [15,16]). However, long electrodes are not ideal for focal stimulation of single cells. The flexible bi-electrode configurations within a densely packed array explored here can provide high density single electrode stimulation while at the same time allowing axon avoidance.

These findings suggest that bi-electrode stimulation can improve the performance of next-generation epiretinal prostheses designed to operate at single cell resolution. Practically, a future device would determine the amplitude of current that must be passed through a given electrode to selectively activate a RGC, create such a mapping for all cell-electrode pairs across the array, and implement an electrical stimulation strategy based on these data to optimize artificial vision [42]. For cases in which a cell is not selectively activatable with a single electrode due to inadvertent activation of a distant RGC at its axon, the device would supply the bi-electrode stimulus to minimize the probability of axonal activation. Thus, the strategy presented here has the potential to mitigate a long-standing problem for epiretinal prosthetic devices.

## Acknowledgments

This work was supported by the Stanford Graduate Fellowship (RSV), Research to Prevent Blindness Stein Innovation Award, Wu Tsai Neurosciences Institute Big Ideas, NIH NEI R01-EY021271, and NIH NEI P30-EY019005 (EJC), F30EY030776-03 (SSM), Polish National Science Centre Grant DEC-2013/10/M/NZ4/00268 (PH), AGH UST, task No. 11.11.220.01/4 within subsidy of the Ministry of Science and Higher Education (WD). We thank Jose Carmena, Mike Taffe, and the California National Primate Research Center for access to primate retinas, and Nishal Shah for useful discussions.

